# Is tumor mutational burden predictive of response to immunotherapy?

**DOI:** 10.1101/2020.09.03.260265

**Authors:** Carino Gurjao, Dina Tsukrov, Maxim Imakaev, Lovelace J. Luquette, Leonid A. Mirny

## Abstract

Cancer immunotherapy by checkpoint blockade (ICB) is effective for various cancer types, yet its clinical use is encumbered by a high variability of patient response. Several studies have reported that the number of non-synonymous mutations (Tumor Mutational Burden, TMB), can predict patient response to ICB. This belief has become widespread and led to the FDA approval of immunotherapy patient prioritization based on TMB levels. The notion that TMB is predictive of response to immunotherapy is rooted in the neoantigen theory. It stipulates that cancer-specific mutations can form neoantigens recognized by the immune system; the more mutations a tumor has, the more likely the immune response is triggered. Here we revisit the data underlying the reported association of TMB with response, and the neoantigen theory. First we assembled the largest pan-cancer dataset of immunotherapy patients with sequencing and clinical data. Surprisingly, we find little evidence that TMB is predictive of response to ICB. We demonstrate that associations similar to the ones reported previously can be observed in shuffled data, suggesting that previous studies suffered from a lack of correction for multiple hypotheses testing and confounding disease subtypes.

Second, we develop a model that expands the neoantigen theory and can be consistent with both immunogenicity of neoantigens and the lack of association between TMB and response. Our analysis shows that the use of TMB in clinical practice is not supported by available data and can deprive patients of treatment to which they are likely to respond.

## Introduction

Immune checkpoint blockade (ICB) treatments such as anti-CTLA-4 and anti-PD1, which target regulatory pathways in T-lymphocytes to enhance anti-tumour immune responses, have already proven to elicit durable clinical responses for some patients (1–4). However, the genetic determinants of response to immunotherapy have yet to be found. Several studies (5–9) suggested that Tumour Mutational Burden (TMB), computed as the total number of nonsynonymous somatic mutations, is correlated with response to immunotherapy in cancer. The underlying hypothesis posits that a fraction of nonsynonymous mutations become exposed as epitopes and constitute neoantigens, which can trigger an anticancer response by the immune system. The association between high mutational burden and response to immunotherapy, within and across cancer types (10–14), has been widely reported in the scientific literature and the media. As a result, TMB is currently discussed as the most clinically advanced biomarker of response to immune checkpoint blockade (15, 16), and the FDA approved the use of TMB to identify patients most likely to derive clinical benefit (17). These studies also triggered a search for inexpensive assays to evaluate TMB directly (18), as well as TMB-derived measures, such as neoantigens, neoepitopes, and mutation clonality (19), which are all currently under investigation to further stratify patients most likely to respond to immunotherapy. Our analysis focuses on TMB itself, as this is the most widely used and only FDA-approved measure.

Using a biomarker to stratify and prioritize patients for treatment runs a risk of depriving patients who have a chance to respond to a life-saving treatment. High variability of response makes relying on a predictor particularly risky. Hence, we revisit original data that were used to establish correlation between TMB and response. We tested TMB as a predictor of both binary responder/non-responder labels from original clinical studies, as well as continuous survival data. We also investigated whether a TMB threshold could distinguish patients with high and low survival after multiple hypothesis testing. We find that no TMB threshold performs better on the clinical data than on randomized ones. We further show that irrespective of the strategy to choose the threshold, even if we were to employ the optimal TMB cutoff, it would still lead to about 25% of responders falling below the treatment prioritization threshold. In addition, we re-examine the pan-cancer association of TMB with response rate to ICB.

Finally we revisit the neoantigen theory that was the rationale for using TMB as a predictor of response to immunotherapy. The theory stipulates that non-synonymous mutations can lead to the production of unique antigens (*neo*antigens) that are recognized by the immune system as foreign, triggering the immune response against cancer cells. The theory further assumes that the more mutations a cancer has, the more likely it triggers the immune system, and the more likely it will benefit from immunotherapy. We develop a simple model that is based on the neoantigen theory and find that it has two regimes. In one regime, the probability of response increases gradually with TMB, as commonly believed. Yet in the other, the probability of response saturates after a few mutations, making a chance to respond independent of TMB. Our analysis of the clinical data is consistent with the latter regime. Thus our model shows that the neoantigen theory is fully consistent with the lack of association between TMB and response.

## Results

### Data aggregation

To evaluate the association of TMB with response to ICB across a broader range of cancer types, we aggregated and analysed data for 882 immunotherapy patients with publicly available pre-treatment whole-exome sequencing data (referred below as CPI800+, **Table S1** and **Material and Methods**). We included patient-level data from an aggregate of early seminal studies (20) as well as a clear cell renal cell cancer (21, 22), non-small cell lung cancer (9), bladder cancer (23) and melanoma (5, 24, 25) ICB-treated cohorts. For every dataset examined, we retrieved TMB levels and survival data (Progression-Free Survival (PFS) or Overall Survival (OS)) for each patient. The original studies provided response classification for most patients.

We also leveraged 1283 patients (termed CPI1000+) who underwent immunotherapy (26), have unified TMB (n=1083) and response definition, as well as survival measures for some patients (n=545 with OS data). Furthermore, we obtained gene panel data (MSK-IMPACT) for 1662 patients (**Table S1**) who underwent immunotherapy. To the best of our knowledge, together this dataset constitutes the largest pan-cancer aggregate of ICB-treated patients with sequencing and clinical data, which allow a robust unified statistical assessment of TMB as a predictor of ICB response

### Is TMB associated with response after treatment?

First, we compared the TMB in patients that have been classified as responders and non-responders based on a number of clinical characteristics. All datasets show not only a considerable overlap in TMB between responders and non-responders, but the lack of considerably elevated TMB for responders and a large range of TMB values in each group.

Consistent with previous studies we find no difference in TMB between responders and nonresponders for major cancer types (clear cell Renal Cell Carcinoma, Head and Neck Squamous Cell Carcinoma and Breast Cancer), with only melanoma (mel1 and mel2, p=0.026 p=0.041) and non-small cell lung cancer datasets (lung1 and lung2, p=8.3×10^-6^ and p=7.7×10^-3^) yielding significant differences as reported earlier (9, 20, 21, 24, 27) (**Figure 1 and Figure S1A**). Analysis in CPI1000+ revealed similar results, yet colorectal cancer, and bladder cancers showed significance of elevated TMB for responders (p=0.044 and p=5.4×10^-7^, respectively) (**Figure S1B**). We also found, consistent with previous studies, that this small elevation of TMB for responders is due to confounding effects of cancer subtypes, i.e. due to the different response rates of cancer subtypes with different TMB ranges (see **Supplemental Text)**. When we revisit a study that reported pan-cancer correlation between response rate and TMB (12, 14), we found that association was driven largely by overall high response in melanoma and subtypes of colorectal cancer (MSI+) with extreme differential response (see **Supplemental Text**). No association between TMB and response rate across all other cancer types is present in available data.

**Figure 1:**
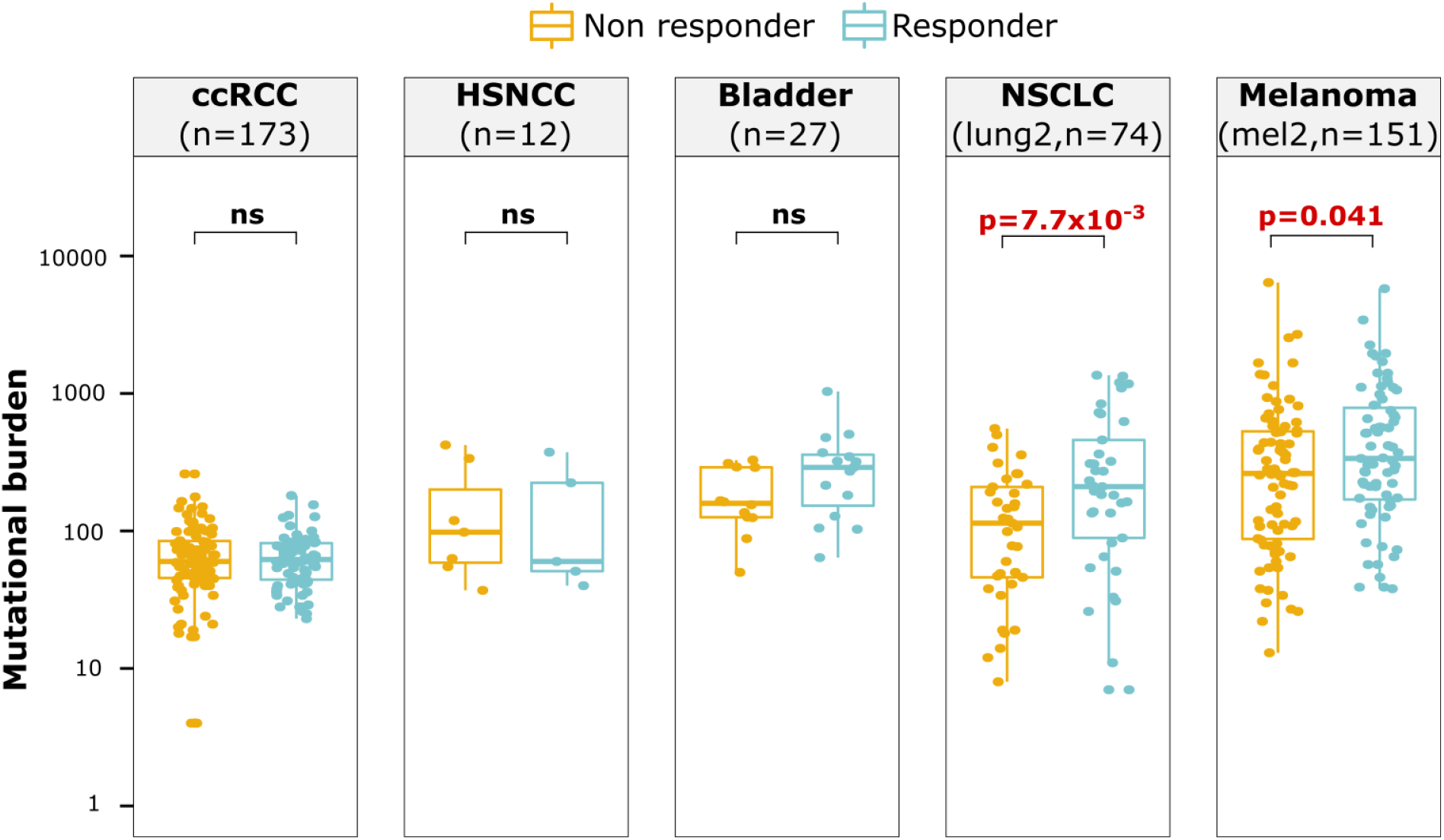
TMB and response to ICB. Association of TMB with response to ICB across five cancer types from CPI800+ (the largest cohorts of each cancer type are plotted here, the others are shown in **Figure S1A**). Only melanoma and non-small cell lung cancer have a significantly different TMB between responders and non-responders. ccRCC: clear cell Renal Cell Carcinoma, HNSCC: Head and Neck Squamous Cell Carcinoma; NSCLC: Non Small Cell Lung Carcinoma

We also examined potential correlations between TMB and survival, rather than using a binary response variable (**Figure 2**). Strikingly, plots of survival versus TMB do not show a visible correlation, trend or TMB cutoff that could differentiate longer and shorter surviving patients. There is a considerable and non-diminishing fraction of patients with long survival, even for lower ranges of TMB values. As we demonstrate below, attempts to find a TMB cutoff value to differentiate long- and short-survivals on such data can be prone to misinterpretation and require careful correction for multiple hypothesis testing.

**Figure 2:**
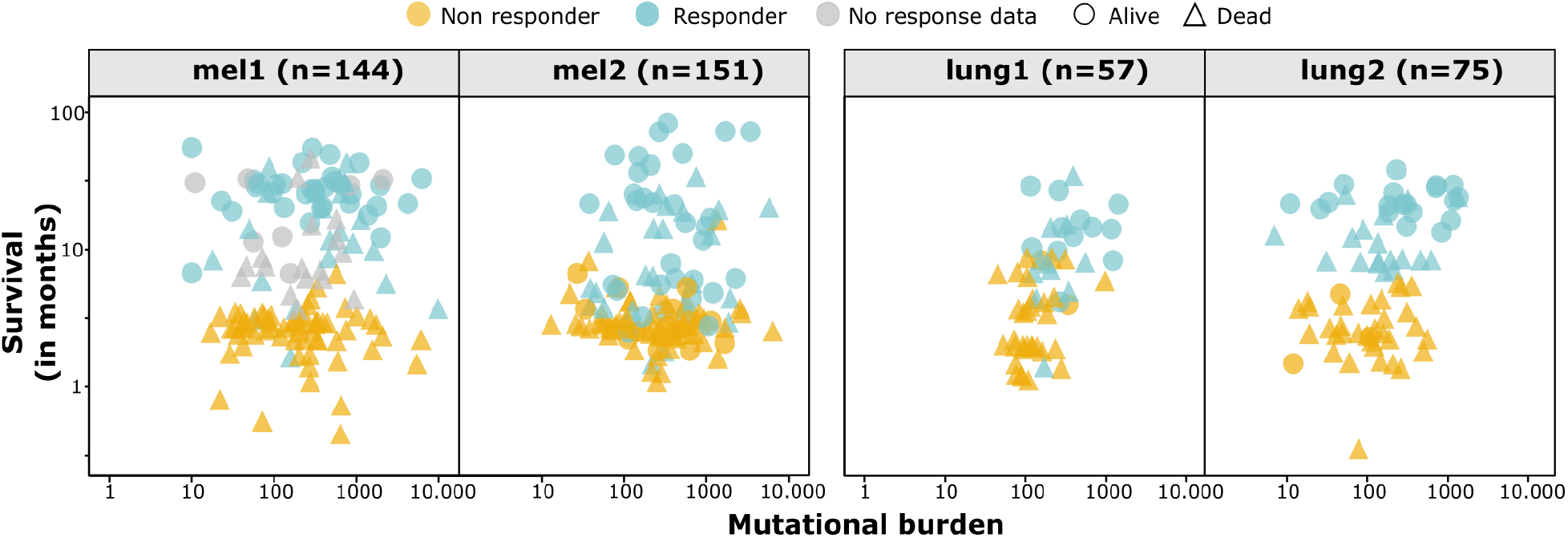
TMB association with progression-free survival post-immunotherapy. Plots of progression-free survival and TMB for melanoma and lung cancer ICB cohorts show the lack of correlation or of an obvious TMB cutoff. Computing a simple correlation for survival and censored data cannot correctly represent the dependence since patients who are alive live longer than the reported survival, and limiting correlation to patients who are dead would bias the analysis. Thus other survival statistics are used through the paper.

Next we investigated whether (i) small differences in TMB between responders and non-responders for some cancer types can be of clinical use, even when p-value>0.05; (ii) there is a potential TMB cutoff that can predict groups of patients with a significant difference in survival; (iii) these clinical data can be reconciled with the neoantigen theory.

### TMB is a poor predictor of response

The key component for validating a biomarker is acceptable classification accuracy, i.e. the biomarker’s capacity to correctly classify a patient’s response (28). ROC curves analysis (**Figure 3A** and **3B**) is a standard tool used across disciplines for measuring the quality of a predictor; it provides a comprehensive quantification of specificity and sensitivity over all possible cutoffs, with the Area Under the ROC Curve (AUC) being an aggregate measure of predictor performance (AUC=0.5 for a predictor performing as well as random). Our ROC-curve analysis shows the (i) lack of a clear TMB cutoff that could be used in the clinic; (ii) poor performance of the TMB-based predictor of response to ICB, as evident from the low AUC in most datasets: mel1 and mel2 yielding of 0.62 and 0.59, and lung2 has an AUC of 0.68. Lung1, however, has the highest AUC of 0.85, which, as we show below, is still insufficient to select patients for ICB. ROC curve analysis on CPI1000+ cohort, with unified TMB, also shows a similarly poor AUC of 0.6 (**Figure S2**).

**Figure 3:**
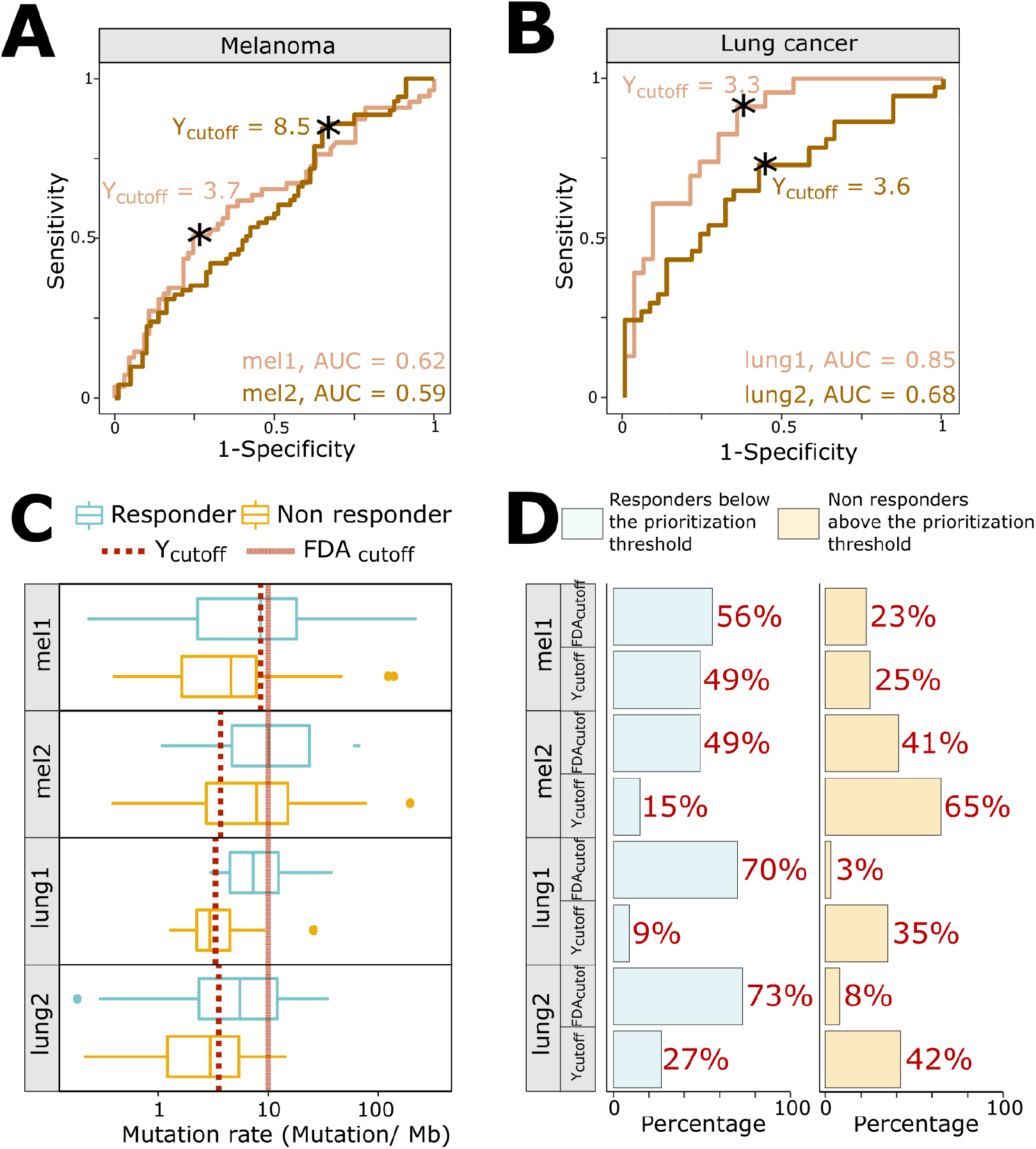
TMB as a biomarker of response to immunotherapy. **(A)(B)** ROC curves for the melanoma and lung cancer cohorts. The asterisk represents the Youden index cutoff. **(C)** Boxplots of nonsynonymous mutation rates across responders and non responders in the melanoma and lung cancer cohorts. The FDA-approved cutoff (10 mutations/Mb) and the best cutoff (Youden index associated cutoff) are shown by vertical lines. **(D)** Proportion of misclassified patients based on the FDA-approved cutoff, as well as the Youden index cutoff for each dataset. The use of either cutoff leads to substantial fraction of misclassified patients (potential responders below the treatment cutoff, or non-responders above the cutoff).

Using a poor predictor for treatment decisions can lead to patient misclassification, i.e. patients who could benefit from the therapy would be deprived of it (responders below the TMB threshold), and patients who get the treatment but don’t benefit from it (non-responders above the threshold). To quantify the shortcomings of TMB-based selection of patients for treatment we computed the proportion of misclassified patients based on the FDA approval of 10 mutations/Mb threshold to select patients for ICB (**Figure 3C** and **Figure 3D**). We find that, on average across the non-small cell lung cancer and melanoma datasets, 62% of responders were below the treatment prioritization threshold and 19% of non-responders were above. While these misclassification rates vary across datasets, fractions of potential responders under the TMB threshold remain high. Moreover, the poor predictive power of TMB indicates that current efforts of choosing a single TMB measure for all cancer types (so call “harmonizing” TMB) would not address fundamental limitations of TMB as the biomarker of response. Indeed, our ROC analysis shows that even the optimal cutoff (Youden index associated cutoff) for each dataset would result in approximately 25% of responders ending up below the treatment prioritization threshold and thus discouraged from receiving a potentially efficacious and life-extending treatment (**Figure 3D**). As such, the main challenge in using TMB in the clinic is the inherently poor association between TMB and response to treatment.

### TMB cannot detect groups of patients with different survival post-immunotherapy

To evaluate the use of TMB for prioritizing patients, and to go beyond the binary response classification, we examined an association between TMB and survival time (Overall Survival, OS and Progression-free Survival PFS). Since survival data is “censored” i.e. only lower bound on survival is known for some patients that didn’t show progression or dropped out of the study, standard correlation-based methods cannot be used to evaluate such association. Nevertheless, groups of patients can be compared using survival analysis methods. Hence we tested whether it is possible to find a TMB threshold that can separate patients into groups with significantly distinct survival.

Scatterplots of survival versus TMB (**Figure 2**, **Figure S3A** and **S3B**) show no evidence of such TMB threshold. Nevertheless, several studies reported (6, 8) a seemingly statistically significant difference in survival between patients below and above a particular threshold. One caveat of this approach is that choosing the threshold value suffers from inherent multiple hypothesis testing, i.e. when the TMB thresholds have been selected among numerous possible alternatives multiple hypotheses are being tested. This inherent multiple hypothesis testing would require further correction of the p-values; a step that is missing in other studies. However, standard approaches (e.g., Bonferroni correction, FDR correction) for multiple hypotheses testing could be too stringent because the hypotheses generated by comparing survival in two groups at multiple TMB thresholds are not independent.

To address this challenge we used a randomization approach (29, 30), similar to earlier studies that examined cutoff selection for dose-response in epidemiological studies (31). We generate 1000 randomized datasets by shuffling TMB among patients, while keeping survival and censored labels unchanged for each patient. First, for each dataset (real or randomized) we determined the optimal threshold that maximizes the difference in survival for groups of patients above and below the threshold. This was done by trying all possible threshold values, computing the difference in survival by logrank test for the groups above and below the threshold; and selecting the optimal threshold that maximizes the difference in survival (i.e. minimize the logrank p-value). Second, we compared the optimal p-value for the real data, *p_real_*, with the distribution of those for shuffled datasets *f(p_shuf_)* and computed the *corrected p-value* as the fraction of shuffled datasets below the real (p_shuf_ < p_real_) (**Figure 4A**).

**Figure 4:**
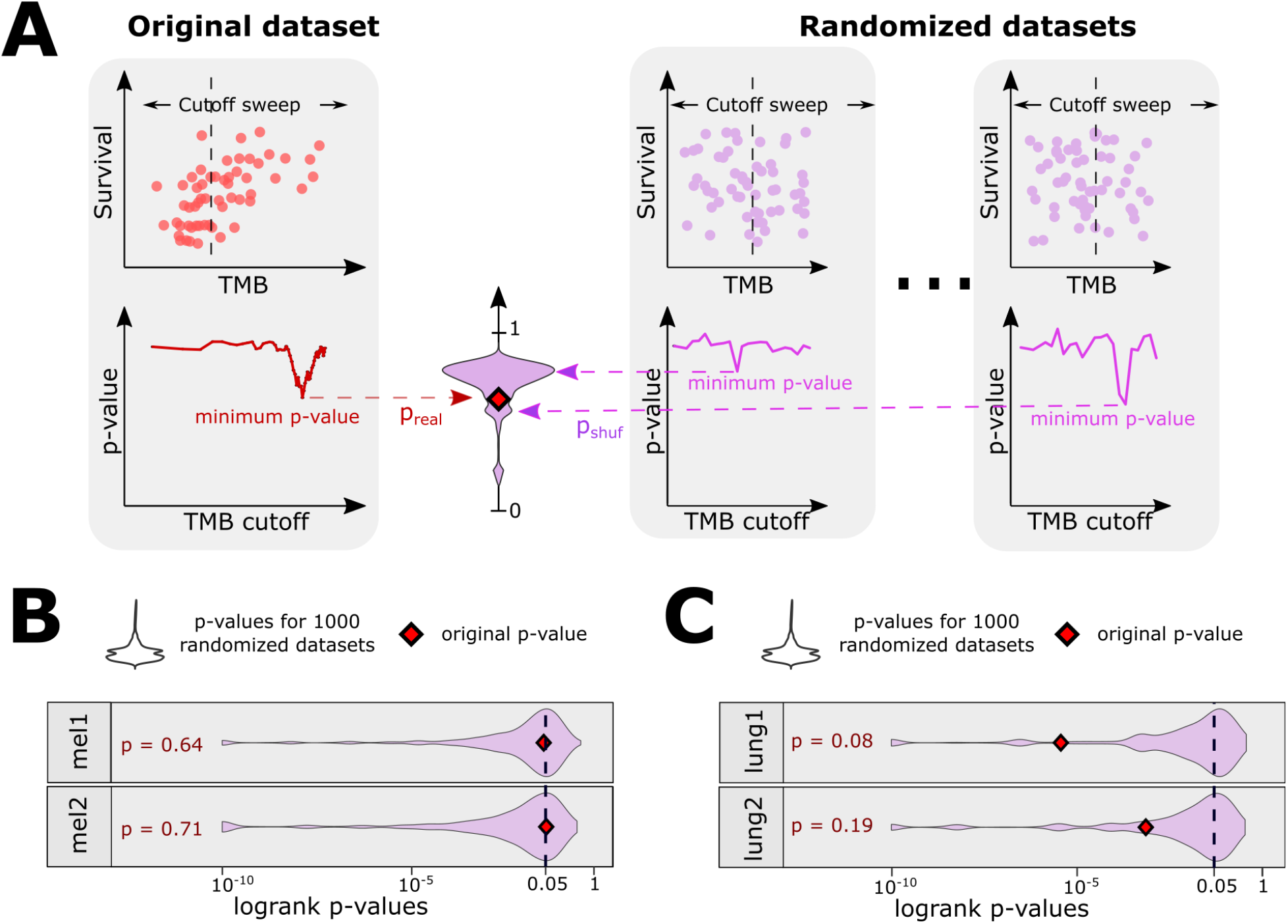
TMB association with progression-free survival post-immunotherapy. **(A)** Overview of the randomization analysis. Left: the optimal cutoff is found to maximize the difference between survival between groups above and below the cutoff (i.e to minimize the logrank p-value, yielding p_real_). Right: the same procedure for shuffled data yields p_shuf_. The fraction of p_shuf_ < p_real_ produces a p-value corrected for multiple hypothesis testing for non-independent tests. **(B)** Results of the randomization analysis in the melanoma cohorts and stratification by subtypes (p-values < 10^-10^ not shown) **(C)** Randomization analysis results in the lung cancer cohorts and stratification by subtypes (p-values < 10^-10^ not shown).

Applied to the melanoma and lung cancer data, this approach (**Figure 4B** and **Figure 4C**) shows that the majority (∼60-70%) of randomly shuffled datasets produced *p_shuf_*below the standard 0.05 threshold, creating a seemingly significant TMB-survival association and emphasizing the need for multiple hypothesis correction. Applied to lung, melanoma and across CPI1000+ dataset (**Figure S4**), our correction for multiple hypothesis testing reveals the lack of a significant TMB threshold that can classify patients into groups with different survival. In particular for lung cancer, for which we previously observed a significant association between TMB and response (**Figure 1A**), we obtained no significant threshold for TMB. We also ran our analysis using OS (for datasets where both OS and PFS are available: mel1, mel2 and lung1) instead of PFS as an endpoint and showed similar results, suggesting that survival definitions do not drive the results of our analysis (**Figure S5A** and **S5B**).

We further obtained consistent results for 1662 patients of MSK-IMPACT cohort treated with ICB but genotyped with gene panels rather than whole-exome sequencing (**Figure S6**). Most of the 10 cancer types tested had a non-significant p-value including colorectal cancer (p=0.088) and melanoma (p=0.093) which have marginally significant p-values, except for non-small cell lung cancer (p=0.034). This study did not provide additional information such as tumor location for melanoma, Microsatellite Instability (MSI) status for colorectal cancer, or COPD for non-small cell lung tumors, which, as we showed above, can confound the association of TMB with response (24, 32). Taken together, our analysis shows the lack of TMB thresholds that can establish a high-TMB group with a significantly longer survival.

### Model reconciles neoantigen theory and data

Neoantigen theory is widely used to argue that cancers with high TMB are more likely to elicit an immune response upon ICB. Although our results show the lack of such dependence, we demonstrate that the effect we observe can nevertheless be explained by a simple mathematical model of neoantigens and immunogenicity (**Figure 5A**).

**Figure 5:**
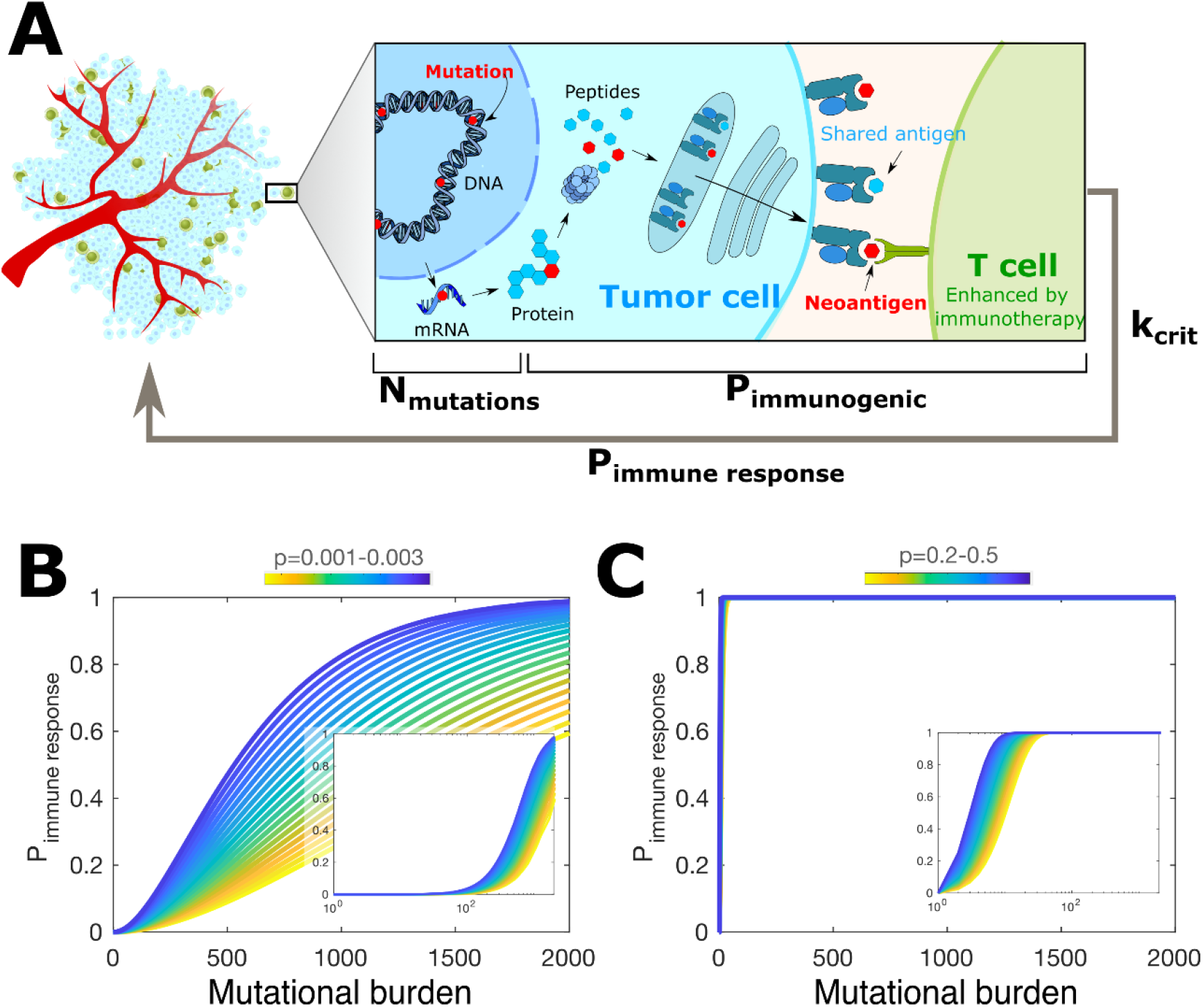
TMB and cancer immunogenicity. **(A)** Our model of cancer immunogenicity coarse-grains several cellular processes into the probability that a mutation becomes immunogenic (*P_immunogenic_*). If the number of immunogenic mutations reaches *k_crit_*, the cancer triggers an immune response. The probability of immune response *P_immune responce_* as a function of TMB for *k_crit_* =2 for low and **(B)** high **(C)** ranges of p, the probability of a mutation to be immunogenic.

Response to ICB treatment requires that cancers elicits an immune response, hence the probability of clinical response can be written as *P*{*clinical response* = *P*{*clinical response*|*clinical response*} · *P*{*immune response*}, where *P*{*clinical response*|*clinical response*} is the probability of clinical response, given that cancer elicits an immune response which is complex and depends on many factors including tumor immune microenvironment. Yet the prerequisite for the clinical response is the immune response *P*{*immune response*} that we focus on. For simplicity assume that every non-synonymous mutation has the same probability *p* to be immunogenic (i.e. to be expressed, presented, interact with MHC, and trigger an immune response; we generalize to different values of *p* below). Due to immunodominance, only *k_crit_* immunogenic mutations are sufficient to elicit a full immune response. Hence, the probability for a cancer with *N* mutations (=TMB) to elicit an immune response is then the probability of having *k_crit_* or more immunogenic mutations among *N*:

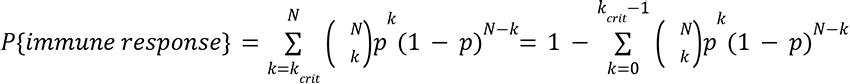

which is the CDF of a binomial distribution. In the case of *k_crit_* = 1, when a single immunogenic mutation can trigger the response the expression simplifies

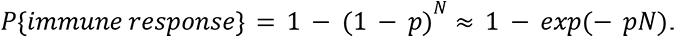

Figure 5B, Figure 5C and **Figure S7** show *P*{*immune response*} as a function of *N* (TMB) for a range of *p* and *k_crit_* values. The model shows two different behaviors. If individual mutations are unlikely to be immunogenic (*p* ≪ 1), e.g. due to a low probability of being presented, the probability of response increases gradually with TMB (Figure 5B). The neoantigen theory generally expects such gradual increase in immunogenicity of cancer with TMB. Yet, available data (Figure 2) don’t show such a trend.

On the contrary, if mutations are more likely to be immunogenic *p*∼0. 1, the probability of response quickly saturates (Figure 5C), making such tumors respond to ICB irrespective of TMB, as we observed in clinical data. In the case of *k_crit_* = 1, *P*{*immune response*} saturates for *N*∼1 and *p* ~ 10. In the data we observe about the same probability of response to ICB for cancers with more than ∼10-20 mutations (Figure 1). Our model shows that achieving about constant *P*{*immune response*} for *N* > 10 − 20 mutations, requires *p* ~ 0. 1 for *k_crit_* = 1, and *p* ~ 0. 2 for *k_crit_* > 1. The same argument holds when each mutation has its own probability to be immunogenic *p*(*i*), then 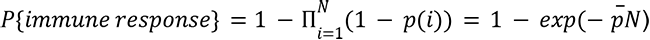, where 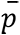 is the mean probability of a mutation to be immunogenic. Thus only the average probability of a mutation to be immunogenic matters. In summary, we find that the model agrees with clinical data if individual non-synonymous mutations have, on average, *p* ~10 − 20% chance for triggering an immune response.

These estimates are consistent with *in silico* neoantigen predictions which showed (33) that 64% of mutations are strong, and 22% weak binders. For these values the model shows that the probability of eliciting a response quickly approaches 1 for TMB ≥ 10 and stays constant and independent of TMB. (**Materials and Methods).**

The model further suggests that for the regime consistent with the data (*P_immunogenic_*=0.2-0.6; *k_crit_* = *1-2*) (i) >90% of tumours with as little as 10 non-synonymous mutations are immunogenic; (ii) when 90% of tumours are immunogenic they have on average as few as 2 immunogenic mutation, the vast majority of mutations present in cancers are non-immunogenic. Such saturation of immunogenicity with low TMB in our model suggests that further immunogenic mutations experience weak negative selection (i.e. threshold epistasis), i.e. weak -if any- immunoediting (34). These results are also consistent with observed immunodominance hierarchies of the T cell responses (35): low TMB tumours can mount responses as robust as high TMB tumours since only a small subset of neoantigens are targeted by T cells.

Furthermore, our model also explains a puzzling observation that immunoediting, i.e. negative selection against immunogenic mutations, is inefficient, allowing tumors to accumulate a high TMB (36, 37). Indeed, once a cancer accumulates mutations making it immunogenic, additional mutations incur at no additional selective disadvantage i.e. show “the epistasis of diminishing return”, and hence accumulate as neutral or weakly damaging passenger mutations (34, 38, 39). Moreover, according to this argument, a cancer should develop means to suppress the immune response early in its development, a prediction that can be tested in future studies of cancer clonal evolution. Taken together, our model and analysis of the available data indicate that cancer with even very few mutations can be immunogenic, suggesting that patients with low TMB might also mount robust immune responses, as has been shown for paediatric patients with acute lymphoblastic leukaemia (35).

## Discussion

Tumor Mutational Burden, a measure of the total somatic nonsynonymous mutations in a tumor, became a popular biomarker of response to ICB, notably because of its relative simplicity to assess.

However, this paradigm is largely based on a series of early papers that examined response in melanoma and lung cancer that we show here to be potentially problematic statistically and further confounded by tumor subtype. Several studies have also reported poor association of TMB with response for specific cancer types, and highlighted TMB and its expression/presentation-based derivatives as problematic for clinical cohort classification (27). In particular for melanoma, published analyses (24) and our results indicate that the site location can explain the observed association between TMB and response to ICB. For lung cancer, our analysis points to the possibility that co-occurrence with COPD may explain the association between TMB and response to ICB among smokers. Overall we demonstrate that while most cohorts and cancer types show the lack of association of TMB and response or survival, the remaining statistical signal in some cohorts can arise due to confounders such as clinical subtypes. Future studies can examine the underlying biology of TMB and neoantigens, aiming to explain the better responses to ICB in certain clinical and cancer subtypes.

Critically, even if responders show significantly but slightly elevated TMB, such associations do not imply the suitability of TMB as a biomarker of response. In particular, we show that no TMB cutoff can distinguish groups of patients with significantly different survival rates. Besides, we show that TMB has poor accuracy as a classifier of response, even in the best-case scenario (Youden optimal cutpoint). This result challenges the FDA approval of TMB for prioritizing patients for ICB. If implemented, such TMB-based clinical decision making would deprive many patients who can benefit from ICB from receiving a life-extending treatment.

An ICB clinical trial that used FDA-approved TMB threshold (KEYNOTE-158) (40) has focused on rare cancers, excluding melanoma and lung cancer. While claiming a higher response rate among high-TMB patients, the trial observed little, if any, difference in overall survival of high-TMB and other patients, putting in question the clinical use of TMB-based prioritization.

We also put forward a simple model that reconciles our findings with the neoantigen theory. Our model shows that if each mutation has a high chance of triggering an immune response, then only a few new mutations make a cancer immunogenic, consistent with the observed immunodominance i.e. the immune response is mounted against only a few of the neoantigens. This result is also consistent with the observed lack of association between antigen density and T-cell presence previously reported (41). Moreover, our model suggests that most cancers are immunogenic, arguing that failures of ICB likely arise due to factors independent of cancer immunogenicity. Quantitative measurements (42) and modelling of neo-antigenic effects can deepen our understanding of cancer development and response to immunotherapy.

Although attractive and scalable, TMB does not consider the effect of specific mutations (missense, frameshift etc), their presentation and clonality (19), nor the state of the tumour, its microenvironment, and interactions with the immune system that can be integrated into potentially better predictors of response to ICB (43, 44). In addition, another major limitation of TMB is the lack of standardized measures. This includes the lack of standard sequencing methods to assess TMB: TMB can be measured from Whole-Exome sequencing, Whole-Genome sequencing, targeted panel and even RNA sequencing(45). This also includes biases introduced by using different mutation calling pipelines resulting in different TMB, sequencing depth and different characteristics of the samples (e.g. low purity samples typically yield lower TMB). For the biology of oncoimmunity, our analysis suggests that, contrary to the neoantigen theory, cancer immunogenicity does not increase with the growing load of neoantigens, and that clinical subtypes can underlie better response to ICB.

Altogether, our analysis indicates that low TMB should not be used to deprive otherwise eligible patients for immunotherapy treatment, and stimulates further research into other determinants of response to immunotherapy.

## Material and Methods

### Immunotherapy study population

CPI800+ was formed of eight independent WES cohorts (n=882, detailed in **Table S1**). The TMB and clinical annotations were not modified from the original studies. Post ICB sequenced samples were excluded from our analysis. In addition, gene panel datasets (n=1662, detailed in **Table S1**) were identified from cbioportal (46).

### TCGA data

Lung cancer TCGA data were also retrieved from cbioportal (46), and additional clinical annotations were downloaded from The Cancer 3′ UTR Atlas (47). COPD status was assessed based on the standard spirometric classification, i.e. post-bronchodilator ratio of forced expiratory volume in one second (FEV1) and forced vital capacity (FVC) below 70%.

## Statistical analysis

We used R version 3.6.2 to perform statistical analyses. Two-group comparisons were evaluated by a two-sided Mann–Whitney U test unless otherwise indicated. P < 0.05 was considered statistically significant.

## Code availability

The R code and data used to reproduce the analysis and figures from the paper are available on GitHub https://github.com/mirnylab/TMB_analysis

## Acknowledgement

We are grateful to Krupa Thakkar, Kevin Litchfield and Charles Swanton for running our analysis on their “CPI1000+” meta-aggregate of immunotherapy patients (26).

We are thankful for many productive and exciting discussions of this work to Christopher McFarland, Johannes Berg, Donate Weghorn, Eli van Allen, Kenneth Kehl, Shamil Sunyaev, Martha Luksza, Michael, Lassig, Boris Reizis, Virginia Savova, Gregory Kryukov, Sebastian Amigorena, Sean McGrath, Baptiste Boisson, Toni Choueiri, Paul B Robbins and all members of the Mirny lab. We are grateful to the organizers and participants of “Physicists working on Cancer” workshop at the Weizmann Institute of Science, Schwartz/Reisman Institute for Theoretical Physics, particularly, Eytan Domany, Herbert Levine, Caterina La Porta and Stefano Zapperi. We are also grateful to the organizers and participants of the workshop “Tumors and Immune Systems: From Theory to Therapy” seminar at Institut d’Etudes Scientifique de Cargèse, particularly, Alexandra Walczak, Thierry Mora, Vassili Soumelis, Paul Thomas and Jason George.

LJL was supported by the Training Program in Bioinformatics and Integrative Genomics (NIH T32HG002295, PI: P.Park). This project grew from the qualifying examination problem in Bioinformatics and Integrative Genomics given by LAM to LJL. We acknowledge support of the MIT-France Seed Fund, and The Chicago Region Physical Science Oncology Center (PS-OC, National Cancer Institute U54CA193419).

---

**Figure S1:**
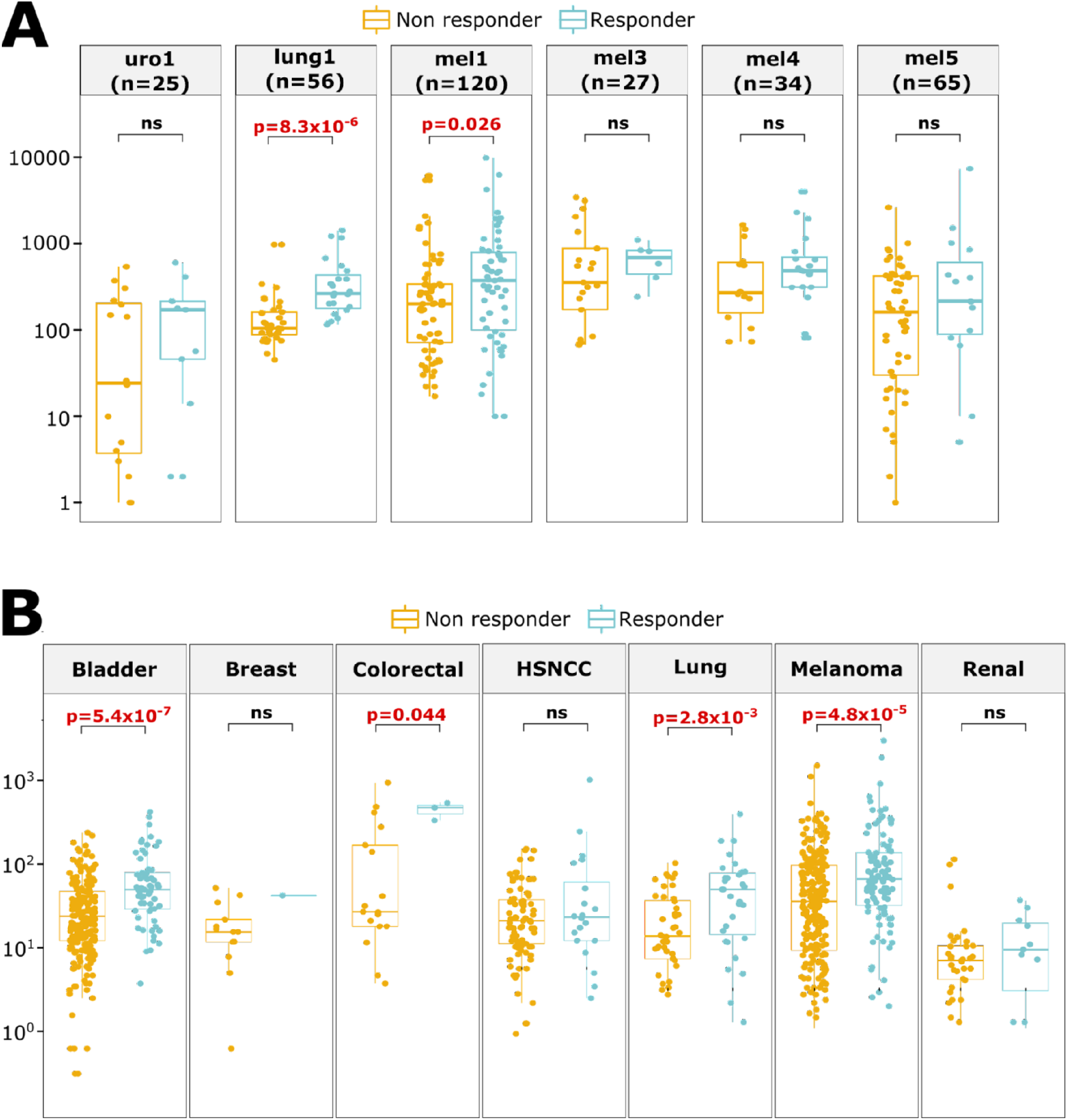
TMB association with clinical benefit from ICB across cancers. **(A)** Association of TMB with response to ICB across three cancer types from CPI800+. Only melanoma and non-small cell lung cancer have a significantly different TMB between responders and non-responders. **(B)** Association of TMB with response to ICB across seven cancer types from CPI1000+. Melanoma, non-small cell lung, bladder and colorectal cancer have a significantly different TMB between responders and non-responders.

**Figure S2:**
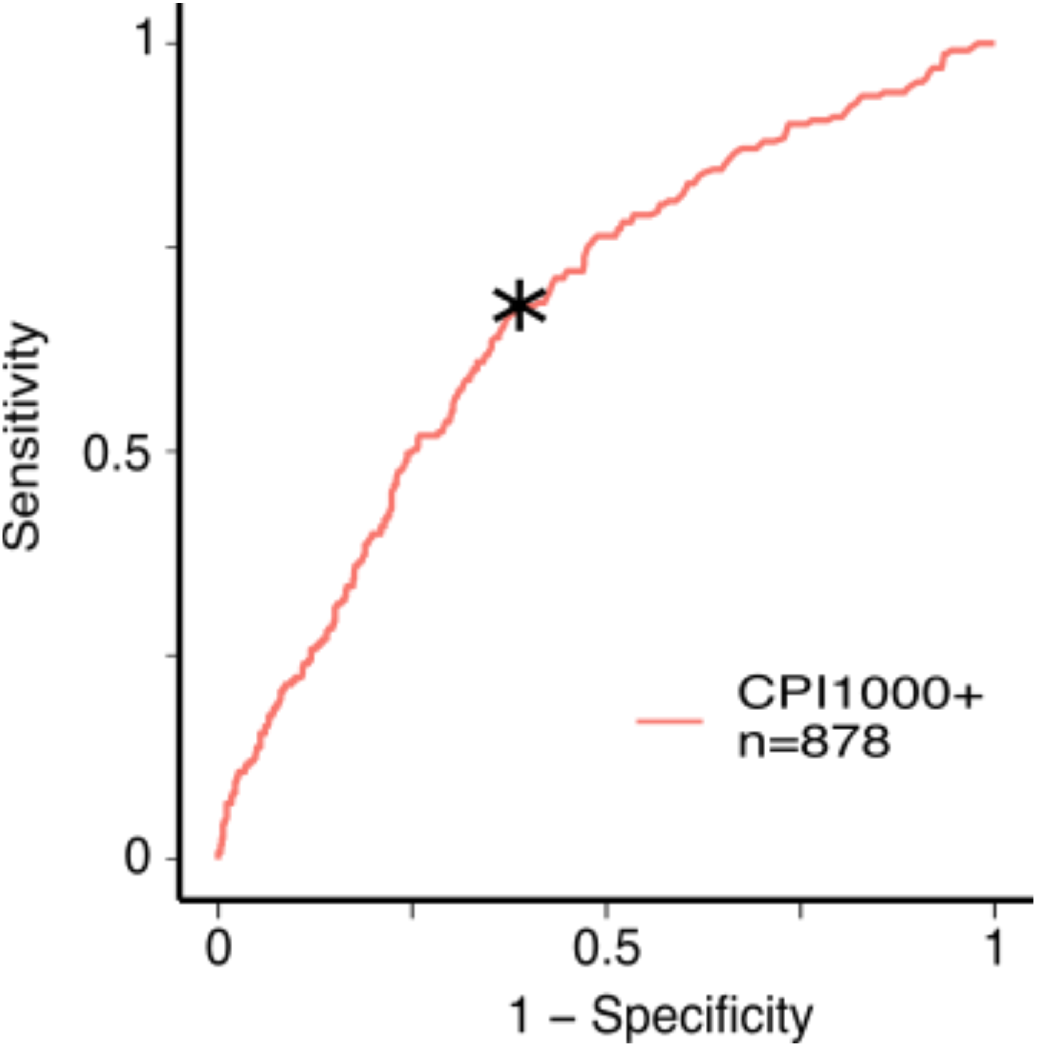
TMB as a biomarker of response to immunotherapy. ROC curve for CPI1000+. The asterisk represents the Youden index cutoff

**Figure S3:**
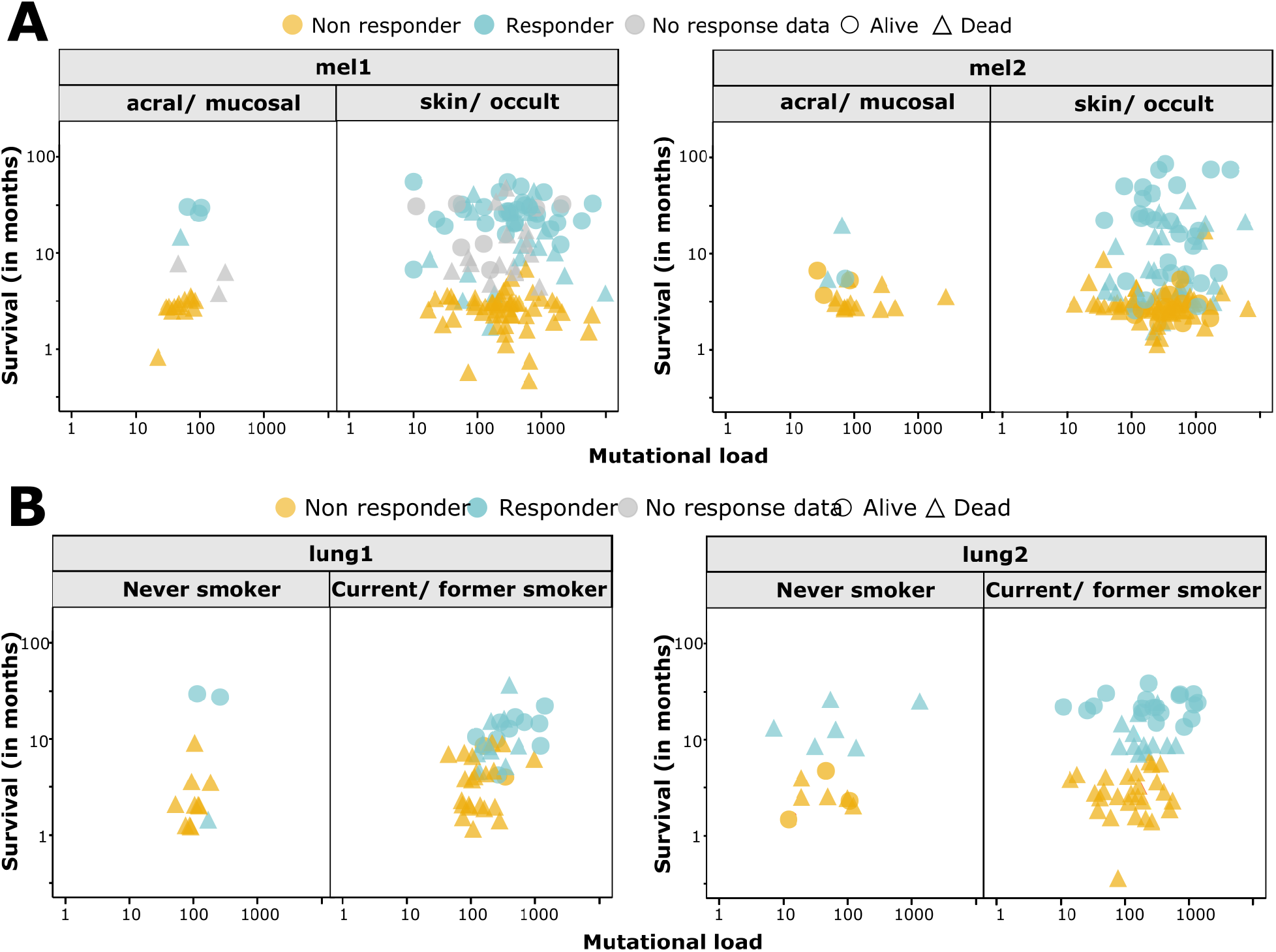
TMB association with progression-free survival post-immunotherapy. **(A) (B)** Plots of progression-free survival and TMB for melanoma and lung cancer ICB cohorts labeled by cancer subtype, showing the lack of correlation or of an obvious TMB cutoff.

**Figure S4:**
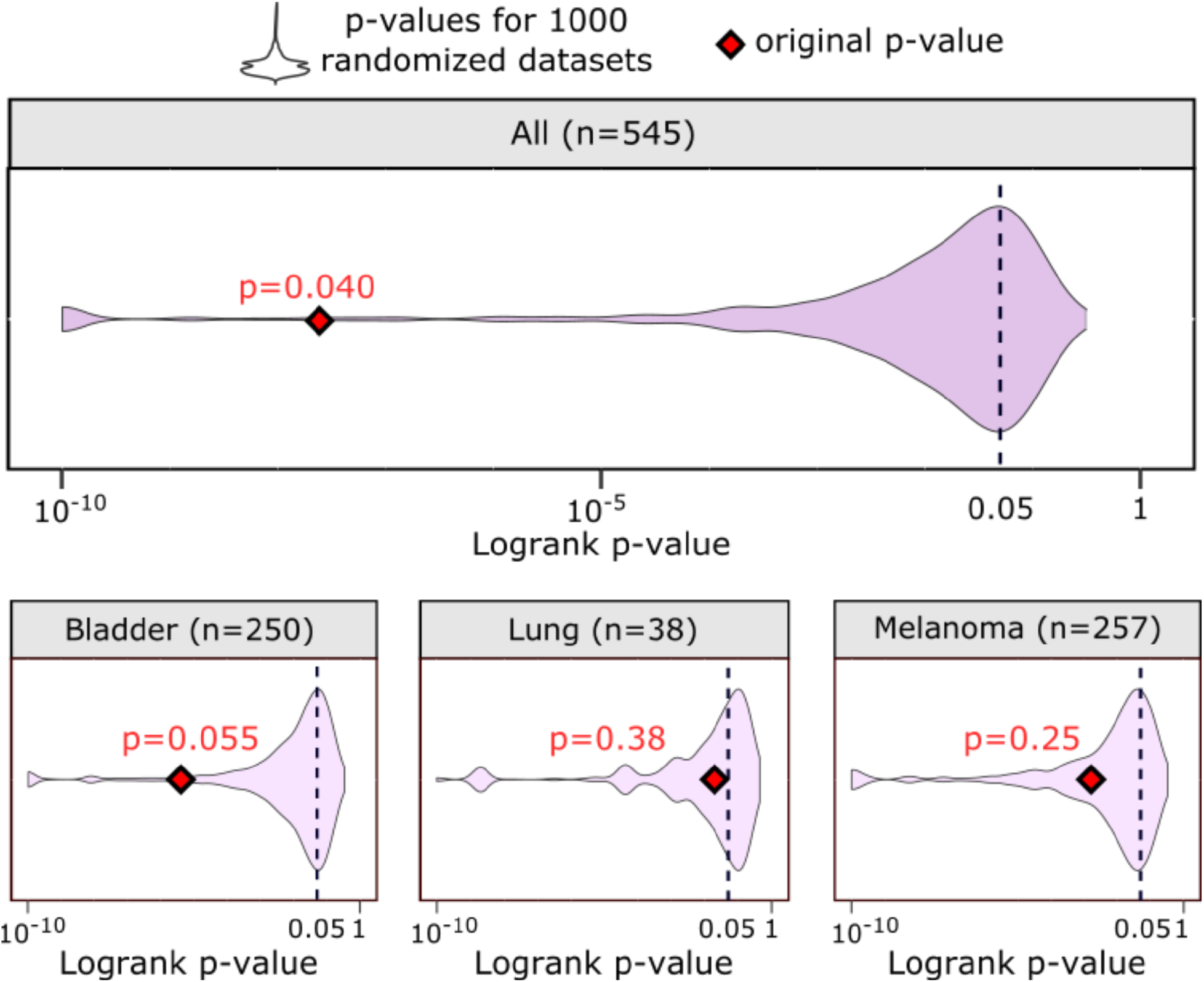
TMB association with overall survival post-immunotherapy. Results of the randomization analysis in CPI1000+ (p-values < 10-10 not shown).When cancer types of CPI1000+ were combined, a nominally significant p-value (p=0.04) arises, likely due to cancer types with different TMB ranges showing significantly different survival rates to ICB.

**Figure S5.**
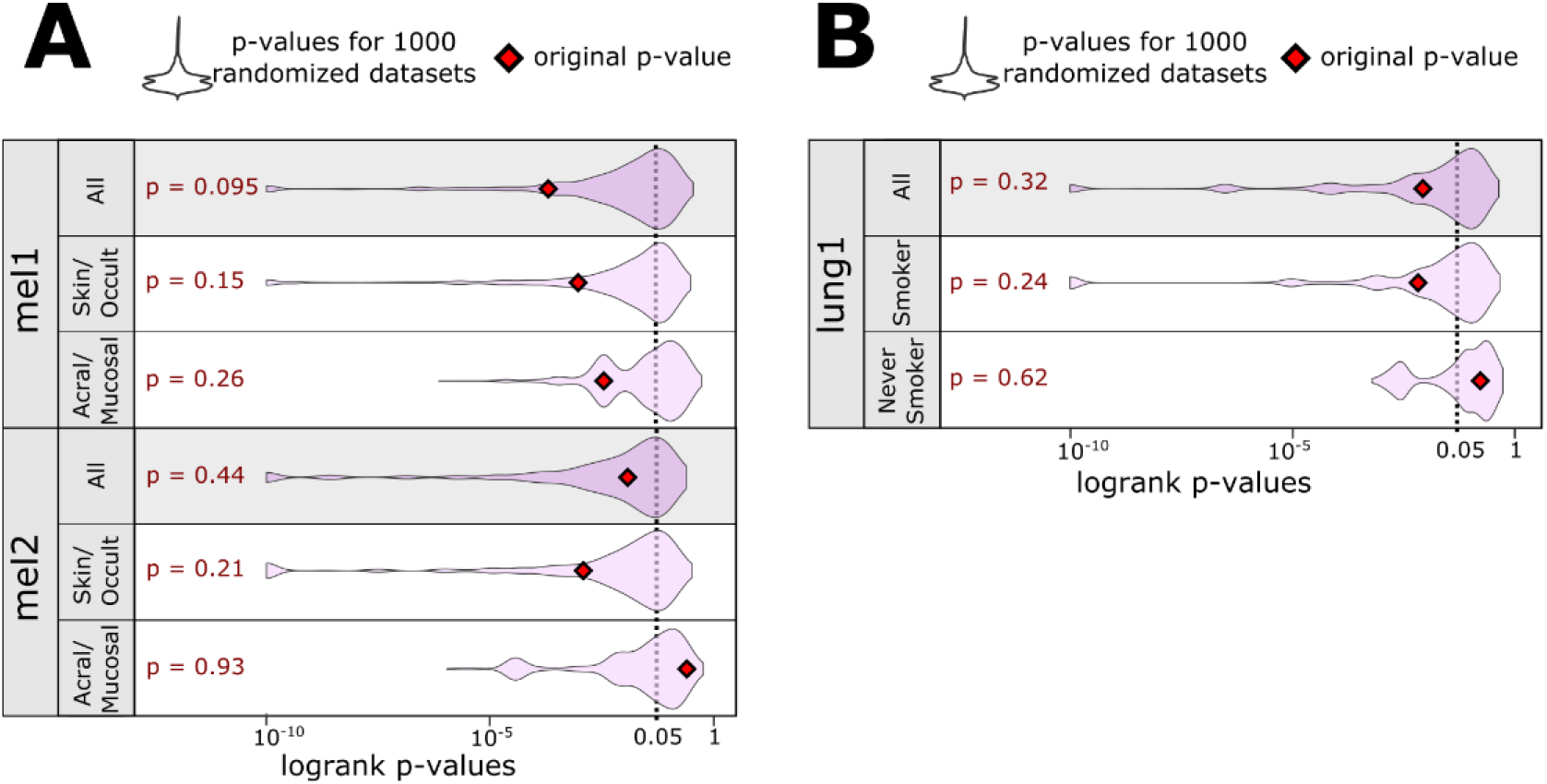
TMB association with overall survival post-immunotherapy. **(A)** Randomization analysis results in mel1 and mel2 and stratification by subtypes (p-values < 10^-10^ not shown) **(B)** Randomization analysis results in lung1 and stratification by subtypes (p-values < 10^-10^ not shown). When corrected for multiple hypotheses all cohorts fail to provide a statistically significant cutoff.

**Figure S6:**
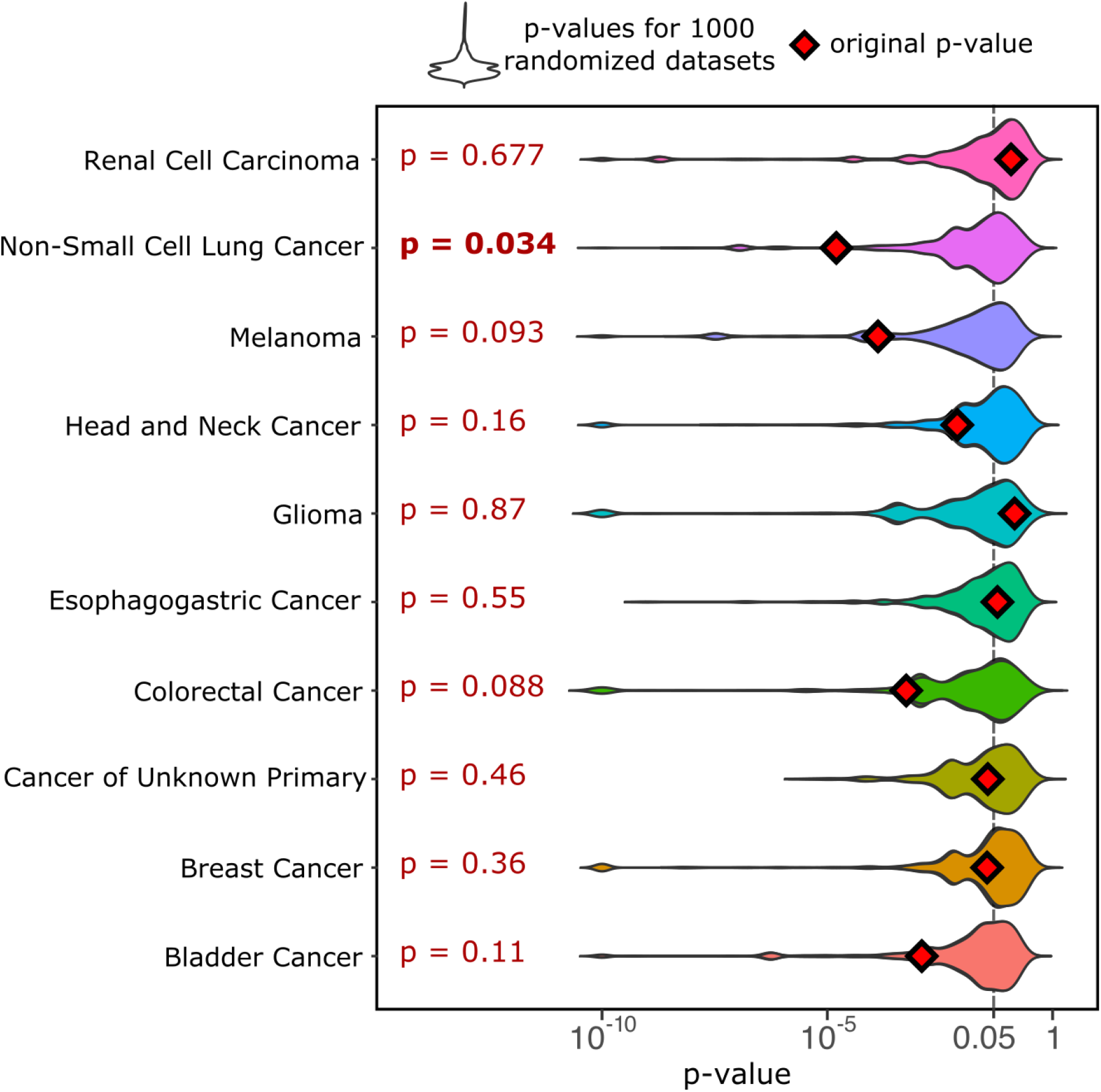
TMB association with overall survival post-immunotherapy. Randomization analysis results in multiple cancer types with MSK-IMPACT targeted next-generation sequencing data (p-values < 10^-10^ not shown)

**Figure S7:**
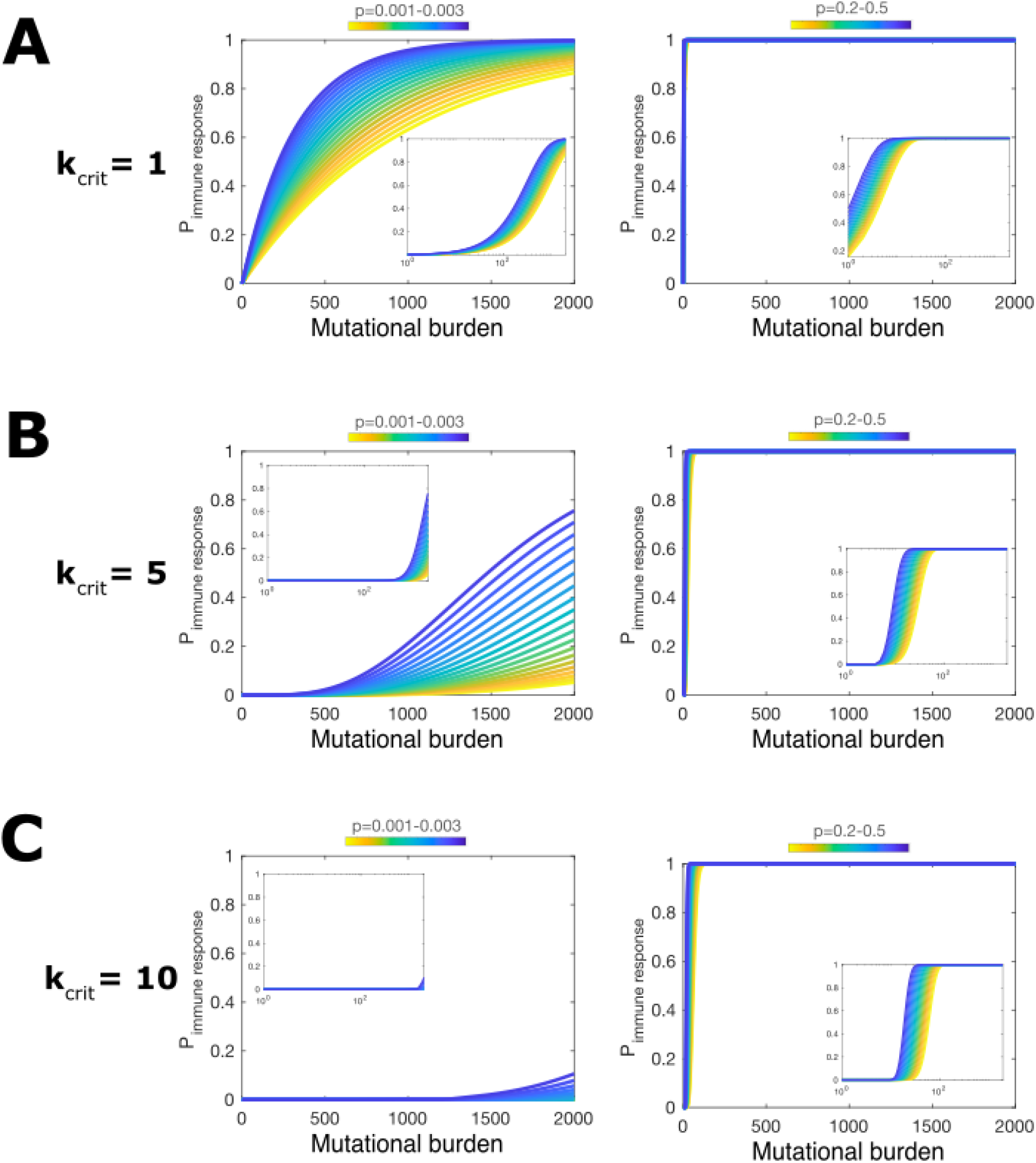
Components of cancer immunogenicity. The probability of immune response *P_immune responce_* as a function of TMB for **(A)** *k_crit_* =1, **(B)** *k_crit_* = 5 and **(C)** *k_crit_* =10 for low (left panels) and high (right panels) ranges of *p*, the probability of a mutation to be immunogenic.

