## Supplemental Table for "Is tumor mutational burden predictive of response to immunotherapy?"

| Cancer type + reference | Dataset naming | Sequencing | Available data | ICB type | Notes |
|---|---|---|---|---|---|
| Various MSS solid tumors [1] | mel2 [1,2,3] (n=151),<br>lung1 [1,4–6] (n=56)<br>HNSCC [1] (n=12)<br>uro2 [1] (n=27)<br>Anal [7] (n=1)<br>SCLC [1] (n=1)<br>Sarcoma [8] (n=1) | WES | Response/ OS/ PFS | anti-PD1<br>anti-CTLA4 | Inclusion criterias defined in [1] |
| Melanoma [9] | mel1 (n=144) | WES | Response/ OS/ PFS | anti-PD1 and or anti-CTLA4<br>(nivolumab/ pembrolizumab with or without prior ipilimumab) | |
| Melanoma [10] | mel3 (n=27) | WES | Response/ OS | anti-CTLA-4 blockade and/ or anti-PD1<br>(pembrolizumab and/or ipilimumab) | 27 patients excluded (21 samples because sequenced on/post ICB + 6 samples because sequenced after the first ICB) NB: when two ICB treatments were given consecutively, we used the OS of the first treatment |
| Melanoma [11] | mel4 (n=34) | WES | Response/ OS | anti-PD1<br>(pembrolizumab or nivolumab) | 4 patients excluded (sequenced on ICB) |
| Melanoma [12] | mel5 (n=68) | WES | Response/ OS | anti-PD1 and or anti-CTLA4<br>(nivolumab with or without prior ipilimumab) | |
| Clear cell renal carcinoma [13] | ccRCC (n=261) | WES | Response/ OS/ PFS | anti-PD1 (nivolumab) | |
| Non-small cell lung cancer [14] | lung2 (n=74) | WES | Response<br>PFS | anti-PD1 &<br>anti-CTLA4<br>(conjoint nivolumab & ipilimumab) | One patient excluded (sequenced post-ICB) |
| Bladder cancer [15] | uro1 (n=25) | WES | Response/ OS/ PFS | Anti PD-L1<br>(Atezolizumab) | |
| | **WES total: 882** | | | | |
| Various cancers [16] | gp1 (n=1662) | Gene Panel | PFS | | |
| | **Gene panel total: 1662** | | | | |

**Table S1: Data aggregate of patients who underwent checkpoint inhibitor immunotherapy.** TMB and response definition was not unified across cohorts
